# Absent from DNA and protein: genomic characterization of nullomers and nullpeptides across functional categories and evolution

**DOI:** 10.1101/2020.03.02.972422

**Authors:** Ilias Georgakopoulos-Soares, Ofer Yizhar Barnea, Ioannis Mouratidis, Martin Hemberg, Nadav Ahituv

## Abstract

Nullomers and nullpeptides are short DNA or amino acid sequences that are absent from a genome or proteome, respectively. One potential cause for their absence could be that they have a detrimental impact on an organism. Here, we identified all possible nullomers and nullpeptides in the genomes and proteomes of over thirty species and show that a significant proportion of these sequences are under negative selection. We assign nullomers to different functional categories (coding sequences, exons, introns, 5’UTR, 3’UTR and promoters) and show that nullomers from coding sequences and promoters are most likely to be selected against. Utilizing variants in the human population, we annotate variant-associated nullomers, highlighting their potential use as DNA ‘fingerprints’. Phylogenetic analyses of nullomers and nullpeptides across evolution shows that they could be used to build phylogenetic trees. Our work provides a catalog of genomic and proteome derived absent k-mers, together with a novel scoring function to determine their potential functional importance. In addition, it shows how these unique sequences could be used as DNA ‘fingerprints’ or for phylogenetic analyses.

## Introduction

Nullomers are short DNA sequences, 18 base pairs (bp) or shorter in length, that do not exist within a certain genome^1^. While the absence of these sequences could be due to mere chance, studies of mammalian genomes have revealed a much larger enrichment of nullomers >10bp than what would be expected by chance^1,2^. One hypothesis that was suggested for their genomic absence, was due to having multiple CpGs which could lead to higher mutation rates^3^. Previous work looking at dinucleotide content has excluded this possibility and suggested that natural selection, i.e. these sequences likely being deleterious to the organism, is a more probable explanation for their absence^2^. This could be due to deleterious properties of a peptide, or if the sequence is noncoding, through an effect on gene regulation, DNA shape, DNA stability or other unknown causes. To date, only three human nullomers have been functionally characterized, two of which were shown to lead to lethality in several cancerous cell types when delivered as exogenic synthetic peptides^4,5^. It has also been shown that some nullomers are conserved between closely related species, for example gorilla, chimp, and mouse^1,2^. A more extreme case of evolutionary exclusion are primes, k-mers that are found absent from all examined species^1^. Previous work utilized the genome of 12 species (human chimp and 10 other non-primates) to identify 60,370 primes of length 14bp or lower.

To identify nullomers that could be deleterious due to protein coding function, a complementary approach could be used; identifying amino acid sequences that are missing from the proteome, termed nullpeptides. Here, we define nullpeptides as sequences that are up to seven amino acids (aa) in length that are absent from the proteome. Previous work carried out in 2009 identified 417 five amino acid primes that are not present in the universal proteome (over 6 million proteins analyzed at that time)^6^. Functional characterization of a 5-mer peptide (KWCEC) that is extremely rare within the universal proteome and absent from the human proteome, showed that it could potentially enhance immunogenicity when administered alongside an antigen^7^.

With the plethora of available genomes, we set out to comprehensively characterize nullomers, nullpeptides and primes in thirty different species. In addition, taking advantage of the annotations available for these genomes, we characterized nullomers in specific functional categories: coding sequence, exons, introns, 5’UTR, 3’UTR and promoters. Using various ranking metrics, we show that nullomers and nullpeptides are under negative selection. Using the large resources available for human variation, such as dbSNP^8^, we characterized how these variants can lead to the materialization of nullomers, termed variant-associated nullomers, and be used as DNA ‘fingerprint’ tools to identify specific individuals. Finally, we show how these sequences could be used to build phylogenetic trees.

## Materials and Methods

### Nullomer and nullpeptide identification

We developed an algorithm in Python that performs an exhaustive search across input sequences, identifying the frequency of each kmer for a selected range of lengths. The algorithm can also be used to identify peptide nullomers using the standard twenty amino acid code and it ignores any rare amino acids such as selenocysteine. Here, we ran the algorithm for nucleotide lengths up to 15 base pairs (bp) and peptide length up to 7 amino acids (aa) from which we derived the list of nullomers and nullpeptides in the reference human genome (hg38) and proteome (UP000005640), respectively.

To assign functional categories in nucleotide nullomers, we utilized the GENCODEv28 annotation^9^ to identify genic and non-genic sequences, with the former being defined as the sequence from the transcription start site (TSS) to the transcription end site (TES). The genic portion was further broken down to 5’UTR, exonic, intronic, 3’UTR and consensus coding sequences (CCDS).

A nullomer of order *i* indicates that any base substitutions in *i* places along the k-mer still result in a nullomeric sequence. Consequently, nullomers of order i+1 are also nullomers of order i. First order nullomers in the human genome and its functional categories were subsequently identified. For each nullomer the algorithm compared the number of occurrences of all possible kmers of one base pair Hamming distance. If the sum of occurrences of all possible kmers across the search space was zero, the nullomer was scored as a first order nullomer.

### Analysis of nullomers across diverse organisms

Nullomer extraction was performed across a range of species using their reference genomes and proteomes. We prioritized all primate species with a good publically available reference genome: chimp, bonobo, gorilla, gibbon, grey mouse lemur, rhesus macaque, Sumatran orangutan, common marmoset, northern greater galago, Philippine tarsier and black-capped squirrel monkey. We added the following additional mammals: pig, horse, cat, dog and cow; and the following rodents: mouse and rat. Finally, we expanded our search beyond mammals to include the house chicken, zebra finch, zebrafish, drosophila (*D. melanogaster*), lizard, nematode (*C. elegans*) and yeast (*S. cerevisiae*). For mouse lemur, a reference proteome was not available and thus was excluded from the nullpeptide analysis. UniProt^10^, which represents a comprehensive and non-redundant database of all known protein sequences across all biological organisms, was used for our nullpeptide and peptide prime analyses. It was downloaded on October 8th 2019 and at the time contained 1,030,456,800 protein sequences.

### Statistical evaluation of nullomers

Expanding on the notion of nullomers of order 1 or higher, we developed a set of statistical methods to prioritize nullomers and as metrics of potential negative selection (Figure 2a). φ1 examines for each possible 1bp substitution of a nullomer *N*_*i*_ the number of times the new sequence occurs. Occurrences for each possible 1bp substitution change of the sequence of *N*_*i*_. Using the median for all possible permutation occurrences yielded φ1. The score was calculated as follows:

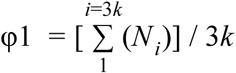

**Equation 1. Score method 1 (SM1) = Expected occurrences.** Median number of occurrences across the one nucleotide variant kmers

**Figure 1.**
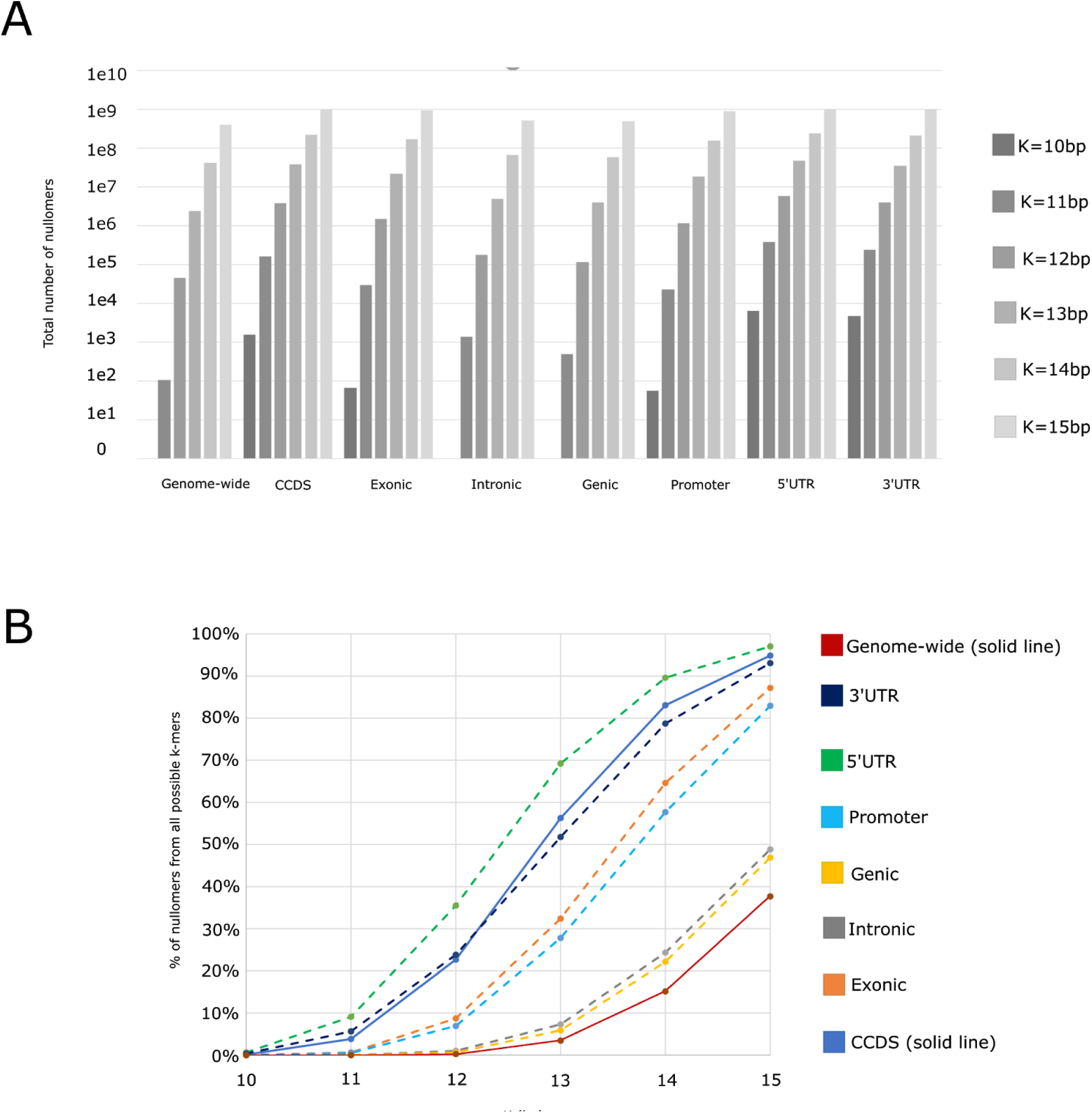
Distribution of nullomers in the human genome and its functional subcompartments. **A.** The number of nullomers in the entire human genome and its sub-categories as a function of nullomer length. **B.** Proportion of kmer space for nullomers K=10-15bp in the human genome and its functional sub-compartments.

**Figure 2.**
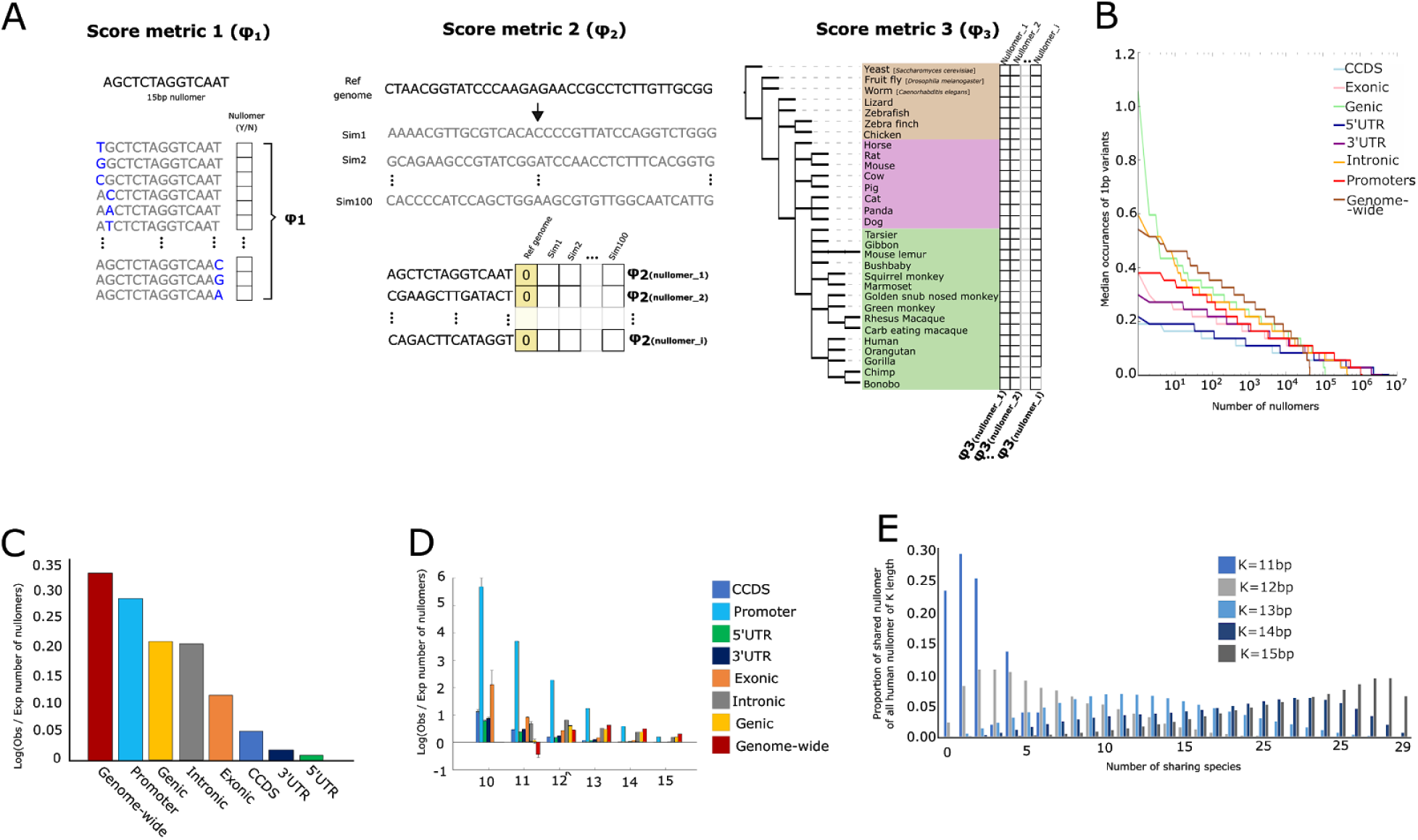
Nullomers are observed more frequently than expected by chance in the human genome and various functional components. **A.** Schematic representation of the various nullomer scoring matrices. **B.** φ1 metric score for each nullomer across the genome and the genomic sub-compartments for K=12. **C.** φ2 metric score associated enrichment of nullomers relative to simulations in the genome and various functional categories across K=10-15 bp. **D.** φ2 metric score associated enrichment of nullomers relative to simulations in the genome and its sub-compartments as a function of nullomer length for K=10-15bp. For **C**-**D**, simulations were performed controlling for trinucleotide content. Error bars represent standard deviation of the number of nullomers obtained from n=100 simulations of each genomic annotation category and has been log transformed. **E.** φ3 metric score showing the proportion of human nullomers found across 29 other species as a function of nullomer length for K=11-15 bp.

A higher value of φ1 might signify that a stronger negative pressure is at play to avoid the nullomer occurrence in the genome. Because the variance in the frequency of each amino acid usage in the proteome is large, for peptides we used a variant of φ1 based on the mean of all possible permutation occurrences of a nullpeptide rather than for all possible 1bp substitutionsS.

Our second score metric (φ2) is based on a 100-fold Monte Carlo simulation permuting the human genome, or its subcompartments, controlling for mononucleotide, dinucleotide or trinucleotide content across each genomic element for each simulation. Simulations were performed using the Ushuffle package^11^. By permuting the genome or its genomic subcompartments, we addressed issues associated with the intrinsic rarity of GC-rich kmers in the human genome. We define φ2 as the average number of times that nullomer *N*_*i*_ appears in our permuted genome sequences.

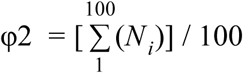

**Equation 2. Score method 2 (SM2) = Expected occurrences.** Averaging the number of nullomer occurrences in 100-fold simulated genomes. A high value of φ2 indicates that the nullomer is potentially under negative selection.

Our third metric added an evolutionary perspective to the score. Strong natural selection against a sequence will be implicated by the absence from the genomes of many species. The species-based nullomer score (φ3) is centered on the occurrence of a k-mer DNA motif in the genome of 30 different species (see above) and it is defined as the average number of species other than human.

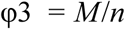

**Equation 3. Score method 3 (SM3) = Occurrences along evolution.** Averaging the number of a species presenting a human k-mer nullomer (*M*) divided by the overall number of species used in the search space (*n*).

Nullomers that have a value of φ3 of 0, are noted as ‘Primes’ as they are absent from all other species examined.

### Population variation and materialization of nullomers

The short human variation annotation file for the human genome was downloaded on February 3rd 2020 from (ftp://ftp.ncbi.nlm.nih.gov/snp/organisms/human_9606/VCF/00-All.vcf.gz). A custom Julia script was used to identify every annotated variant that can result in the creation of a k-mer which was designated as a nullomer based on the reference sequence, for substitutions and single base pair insertions and deletions. Analysis of the trinucleotide context of mutations and the six possible substitution types was also performed. Analysis of the frequency of nullomers materializing due to variants was performed which was further subdivided by the allele frequency of annotated variants. A similar analysis was done for nullpeptides.

### Phylogenetic analyses of nullomers and nullpeptides

Nullomers were identified separately for each species and for each kmer length. The Jaccard index, which is calculated as the size of the intersection of shared nullomers between a pair of species divided by the size of the union of nullomers in the two species, was used as a similarity metric for the construction of phylogenetic trees. Phylogenetic trees were constructed in Python using the package “scipy”^12^ and the functions “dendrogram” and “linkage” using the Ward’s method as a criterion. The package “seaborn” and the function “clustermap” were used to construct the hierarchical clustering dendrograms and the associated heatmaps using default parameters. The same analysis was performed for the construction of phylogenetic trees using sequences.

## Results

### Nullomer annotation

We first set out to generate a comprehensive list of all human DNA nullomers for each kmer length up to 15bp (**Figure 1a**) (see Methods). The shortest nullomers we could find in the human genome were 11bp long (**Table 1** and **Figure 1a**), with a total of 104 nullomers at this length, consistent with previous reports^2^. The number of nullomers grows exponentially with increasing kmer length (**Figure 1a**; **Table 1**). For example, we find ∼40 million nullomers at K=14 and ∼400 million at K=15 in the human genome. Moreover, for K=12 the nullomer space represents only 0.26% of all possible 12-mers, whereas for K=15, the nullomer space represents 37.8% of all possible 15-mers (**Figure 1b**).

**Table 1.** Number of k≤15bp nullomers found in the human genome and in various functional categories.

| Length | Genome | Genic | CCDS | Exonic | Intronic | Promoter | 5'UTR | 3'UTR |
| --- | --- | --- | --- | --- | --- | --- | --- | --- |
| 10 | 0 | 0 | 1,553 | 65 | 0 | 55 | 6,321 | 4,783 |
| 11 | 104 | 480 | 159,400 | 29,908 | 1,342 | 22,390 | 38,0030 | 235,512 |
| 12 | 44,287 | 115,211 | 3,798,220 | 1,459,145 | 179,991 | 1,157,400 | 5,954,005 | 4,008,652 |
| 13 | 2,347,664 | 3,986,232 | 37,728,254 | 21,795,714 | 4,889,886 | 18,632,872 | 46,458,594 | 34,749,772 |
| 14 | 40,798,250 | 59,680,430 | 222,865,418 | 173,484,802 | 65,434,475 | 155,022,567 | 240,489,521 | 211,238,387 |
| 15 | 405,373,474 | 504,426,674 | 1,018,873,404 | 935,392,848 | 523,843,630 | 889,865,066 | 1,042,269,144 | 999,624,368 |

We next characterized nullomers in different functional categories. These include genic regions [mRNA sequence from transcription start site (TSS) to transcription end site (TES)], consensus coding sequences (CCDS), exons (both coding and noncoding), introns, 5’UTR, 3’UTR and promoters (defined as −2500 to +500 around the TSS) (see Methods). Analogous to nullomers within the entire human genome, we did not find genic or intronic nullomers for K< 11. However, we did identify 1,553 CCDS, 65 exonic, 6,321 5’UTR, 4,783 3’UTR and 55 promoter nullomers at a length of K=10. Similar to our nullomer analyses for the entire human genome, we also observed an exponential increase in the number of identified nullomers with increasing kmer length for the various functional categories (**Figure 1a, Table 1**). In addition, when increasing kmer length nullomers represented a larger portion of the kmer space (**Figure 1b**). This repository of human DNA nullomers serves as the basis for all subsequent downstream analyses and comparisons in this article.

### Nullomers are under negative selection

We next set out to test if these sequences are absent due to mere chance or if they are under negative selection. Lacking a conventional statistical measure to determine if a nullomer is missing, we created our own scoring matrix, φ N. The φ N score encompasses three tiers of ranking (**Figure 2a**), described in detail below. All three tiers estimate deviations from the number of the expected occurrences for each kmer motif (not necessarily a nullomer) in order to identify kmers that were relatively enriched or depleted. Our first score metric (φ1), is the median number of occurrences of all the 1bp possible substitution kmers for each nullomer sequence across the search space. The second scoring metric (φ2), is based on simulations of the human genome, its genomic sub-compartments or the human proteome controlling for mono- di- or tri-nucleotide or mono- di- or tri-peptide content composition, respectively. Lastly, our third score metric (φ3) is evolutionary driven, based on nullomers in the genome of 30 species (including humans) across vast evolutionary distances.

We used simulations of each regulatory component to calculate the number of occurrences of nullomers by chance in the human genome and its sub-compartments. We calculated the ratio between the number of observed versus the number of simulated (expected) nullomers and detected a higher number of nullomers than expected by chance (**Figure 2c**). Simulation scores (φ2) showed a significant nullomer enrichment in the human genome for every nullomer length between K=12-15 (**Figure 2c-d**). For K=11, we find more sequences in the simulations than in the genome, likely due to the small number of nullomers (N=104). We observed the most pronounced number of nullomers compared to the simulations for promoters, genic regions and introns (**Figure 2c**). We also performed the same analysis separating the different nullomer lengths, finding that shorter nullomers displayed a higher nullomer enrichment in the genome sub-compartments (**Figure 2d**). This could be the result of a larger kmer space being nullomers with increasing kmer length (**Figure 1b**). Leveraging φ3, the average occurrences in 30 species (**Figure 2e**), we identified a total of 124 genome primes of 13bp length, consisting of only 0.00022% of the human genome nullomer space of the same length, and 2,702,412 of coding human nullomers (0.04%) are actually primes. Combined, our scoring metrics φ N show that nullomers are significantly under selective pressure. In addition, they can also provide ranking criteria for follow-up functional assays on these sequences.

### Higher order nullomers as a ranking criteria

An alternate approach to rank nullomers that are more likely to have a functional consequence is the annotation of higher order nullomers, i.e. sequences where more than one nucleotide change is needed for them to cease being a nullomer. The hypothesis is that if nullomers are under negative selection, more changes would be needed to disrupt them, meaning they will have a higher order. In addition, this approach could be used to rank nullomers, thereby highlighting nullomers under strong negative selection. We mapped the distribution of first order nullomers in the human genome and its functional subcompartments, where one nucleotide substitution is sufficient to disrupt a nullomer (**Table 2**). Examination of nullomers in the entire genome reveals that 14bp nullomers are the shortest first order nullomers (any single base pair substitution does not destroy the nullomer) found and their number increases by three orders of magnitude at 15bp with over 2.5 million first order nullomers (**Table 2**). Focusing on specific functional categories, we find the most first order nullomers in all the lengths analyzed (up to 15bp) within the 5’UTR, possibly due to the strong selection against stop codons, GC content needs and upstream open reading frames (uORFs) that can have a large effect on protein translation^13^. Of note, for CCDS we found 210 12bp first order nullomers which would be of interest to functionally characterize as they correspond to 4 aa peptide sequences. Comparing the number of zero order (at least one nucleotide substitution is sufficient to destroy a nullomer) and first order nullomers reveals large differences in their frequency. For example, there are over 3.78 million CCDS nullomers of length 12bp, but only 210 (0.006%) of which are designated as first order coding nullomers (**Tables 1-2**). In sum, annotating higher order nullomers could provide a potential ranking approach for follow up functional assays.

**Table 2.**
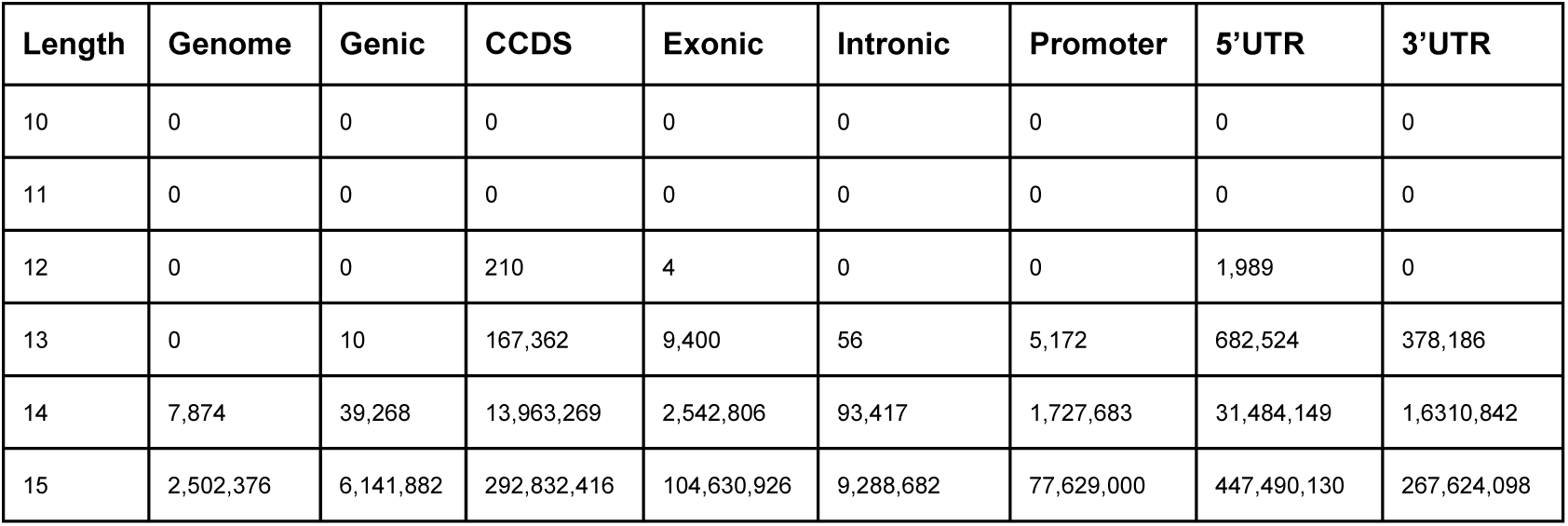
Number of first order nullomers found at each genomic element for kmer up to 15bp.

| Length | Genome | Genic | CCDS | Exonic | Intronic | Promoter | 5'UTR | 3'UTR |
| --- | --- | --- | --- | --- | --- | --- | --- | --- |
| 10 | 0 | 0 | 0 | 0 | 0 | 0 | 0 | 0 |
| 11 | 0 | 0 | 0 | 0 | 0 | 0 | 0 | 0 |
| 12 | 0 | 0 | 210 | 4 | 0 | 0 | 1,989 | 0 |
| 13 | 0 | 10 | 167,362 | 9,400 | 56 | 5,172 | 682,524 | 378,186 |
| 14 | 7,874 | 39,268 | 13,963,269 | 2,542,806 | 93,417 | 1,727,683 | 31,484,149 | 1,6310,842 |
| 15 | 2,502,376 | 6,141,882 | 292,832,416 | 104,630,926 | 9,288,682 | 77,629,000 | 447,490,130 | 267,624,098 |

### Nullomers that materialize due to variation

Nullomers that are absent from the human genome could materialize due to genetic variation. To test this, we generated genome-wide maps of where nullomers could materialize due to nucleotide variation, both nucleotide substitutions and one base pair insertions and deletions. We found 100,587 potential variants that materialize nullomers of 11bp length and 15,085,018 potential variants of 12bp length. The most frequent mutation type was insertions followed by substitutions for both 11bp and 12bp nullomers (**Figure 3a**). The substitutions were further analyzed for their base change. We found that A->C, T->C and G->C are the most frequent substitution types and for trinucleotide context, CTG being the most common both before and after correction for trinucleotide frequency across the genome (**Supplementary Figure 1**). We examined the number of possible mutations that can create each nullomer, finding substantial differences, with up to 25,000 possible mutations generating a subset of nullomers (**Figure 3b**). Throughout the genome we observed that nullomer materializing mutations were more frequent in the genic regions compared to the intergenic (between genes) (**Figure 3c**). Next, we investigated if the putative mutations were more likely to overlap the aforementioned functional categories (genic, CCDS, exons, introns, 5’UTR, 3’UTR, promoters). We found that CCDS, exonic, 5’UTR and promoter regions were the most enriched for variant-associated nullomers, followed by 3’UTR, genic and intronic (**Figure 3c**). The observation that CCDS regions have a higher density of potential nullomer materializing variants, further suggests that these sequences are under strong selective constraints.

**Figure 3.**
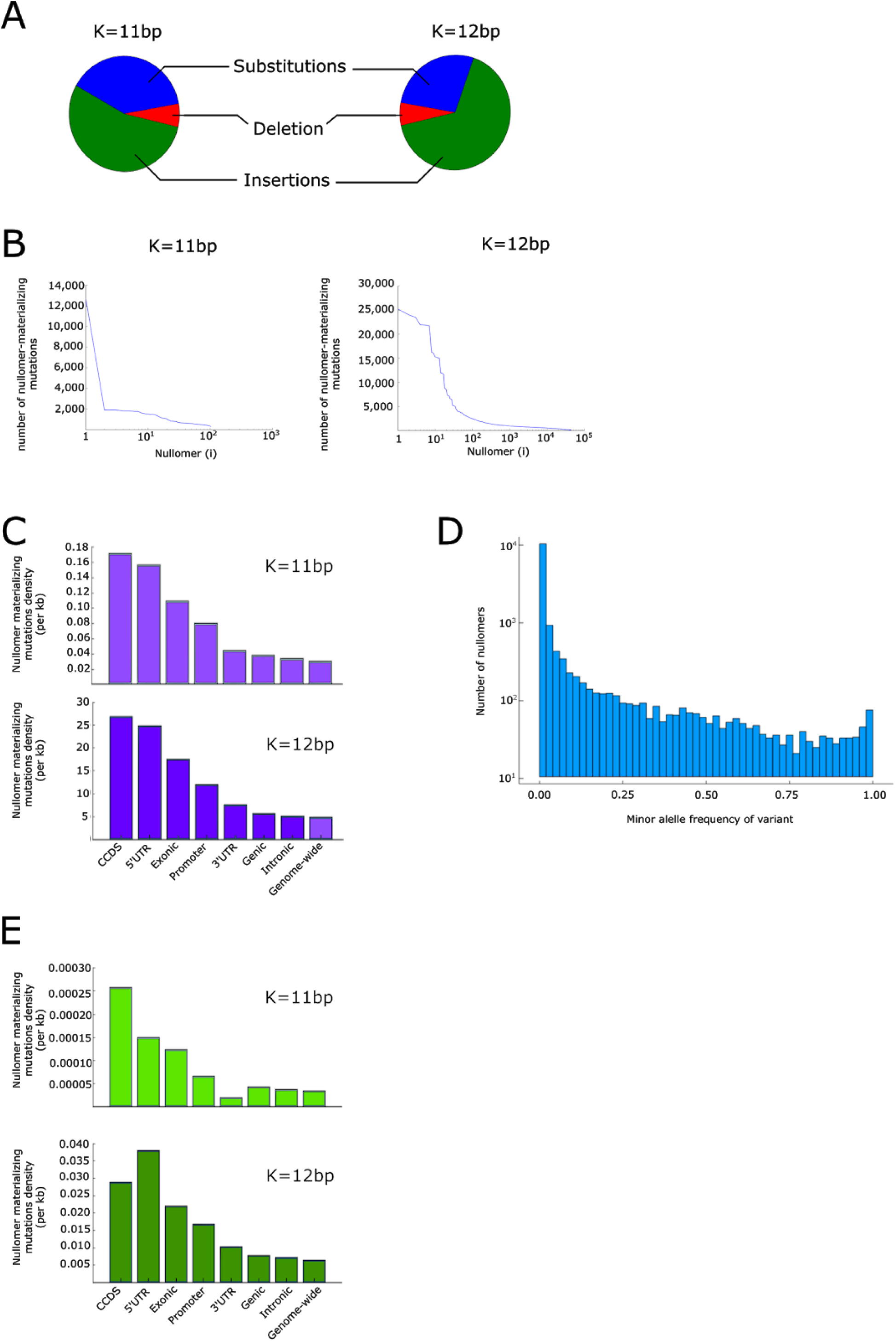
Variant-associated nullomers. **A.** Pie charts representing the proportion of nullomer materializing mutations in the genome that are substitutions and single base pair insertions and deletions for K=11-12bp. **B.** Number of mutations that materialize each nullomer for K=11-12bp across the human genome. **C.** Density of mutations that can materialize nullomers across genomic sub-compartments for K=11-12bp nullomer lengths. **D.** Nullomers can emerge due to population variants. Number of human nullomers per each 2% increments in variant minor allele frequencies. **E.** Distribution of materializing nullomers by allele frequency for K=11 and K=12.

We next set out to test whether nullomers could materialize due to naturally occurring variants in the human population, termed here as variant-associated nullomers. To investigate this, we took advantage of a collection of over 37 million variants annotated in dbSNP^9^. For K=11, we found 107 variants that result in 67 nullomers no longer being absent from the genome. We analyzed the allele frequency of these variants and found that many variants are rare (< 1%). We estimate that 22 of the nullomers have a probability > 5% of being present due to polymorphisms and 33 are present with a probability < 1% (**Figure 3d**). For K=12, there are 18,533 variants that result in 15,815 nullomers no longer being absent. Analysis of the allele frequencies suggests that 3,892 nullomers are likely to be found in at least 5% of the population and 9,489 in < 1%. We next mapped those mutations across functional genomic regions and compared the density of these mutations in the whole genome and in each of them individually (**Figure 3e**). We found that the two most enriched categories for variants that materialize nullomers were 5’UTR and CCDS regions (**Figure 3e**), similar to the findings we observed for the genome-wide maps of potential mutation sites (**Figure 3c**).

### Nullomer annotation across evolution

To identify nullomers that are conserved across evolution and in specific classes, order or species, we repeated our analyses on 29 additional species. These include 14 primate genomes and 15 non-primate genomes (see **Supplementary Table 1** for list of species). In addition, we also annotated nullomers in these genomes using the various functional categories (genic, CCDS, exonic, intronic, 5’UTR, 3’UTR and promoters). We observed a negative correlation between genome size and number of nullomers (**Figure 4a**), (Pearson r = −0.89). We also saw that the longer the nullomer, the larger the proportion was shared between species (**Figure 4b**). For K=12, we did not find any shared nullomers between species.

**Figure 4.**
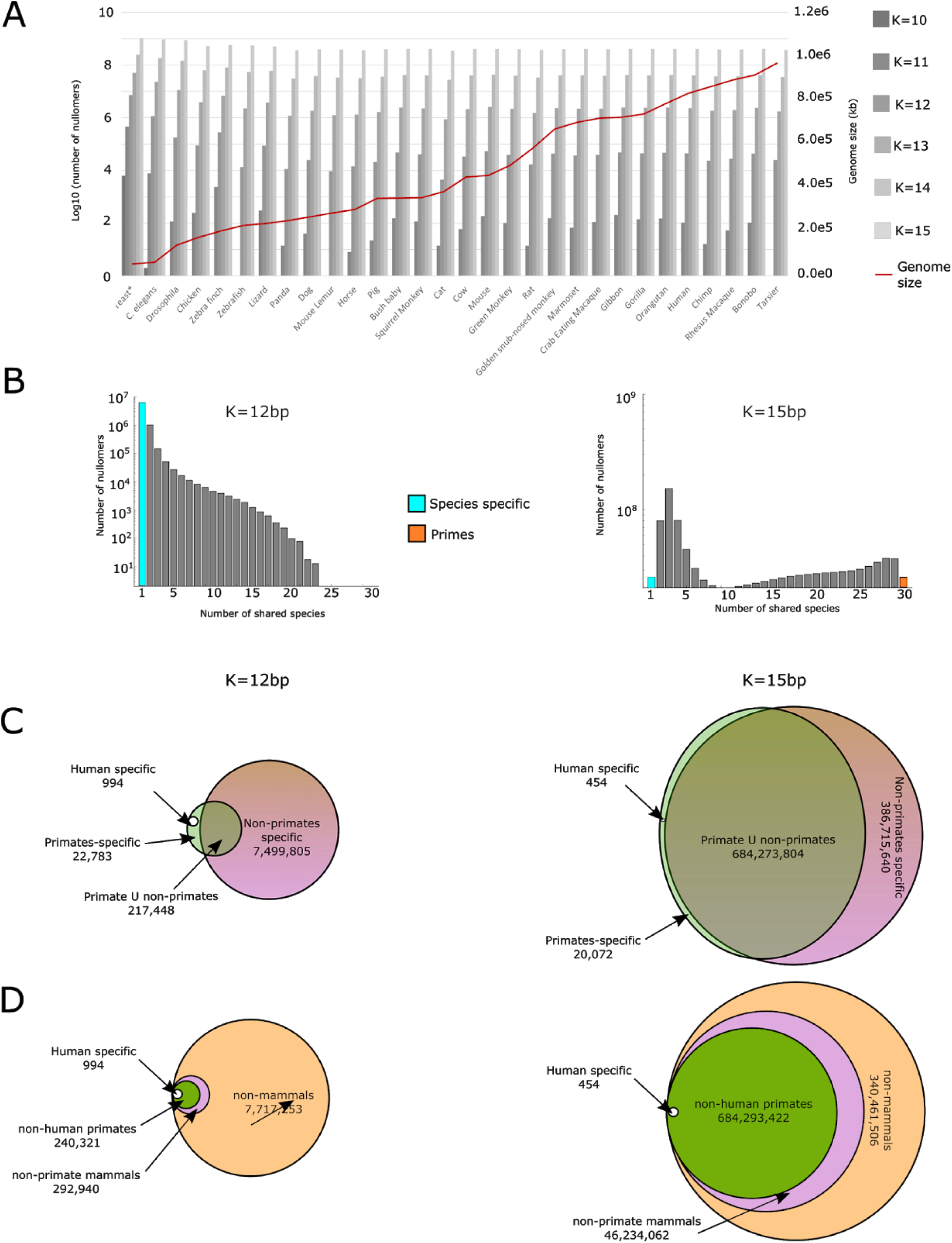
Characterization of nullomers across the genome of 30 species. **A.** A bar plot representing the number of nullomers found per species as a function of kmer length. The red line represents the genome size of each species. **B**. Number of nullomers shared when increasing the number of species for K=12bp and K=15bp. **C**. Intersection of nullomers between humans, all other primates aside from human and non-primates species. **D**. Intersection of nullomers between humans, all other primates aside from human, all mammals beside primates and all species. The diameter of the circles is roughly correlated to differences in numbers.

We next set out to identify nullomers that do not exist in all 30 organisms, termed as nullomer primes, and ones that are unique to each species (**Figure 4b**). We found 124 nullomer primes at length 13bp (0.00022% of all nullomers of similar length), 272,085 primes at length 14bp and 26,010,370 primes at length 15bp (2.4% of all nullomers of similar length). For CCDS, K=12 were the shortest primes we found, with a total of 18,966 K-mers. In terms of organism specific, we compared human nullomers to non-human primate nullomers (14 species), to reveal that although there are no primes for kmers of 12bp, 994 (0.0129%) are human specific nullomers, and a total of 22,783 (0.136%) are primate (including human) specific (**Figure 4c; Supplementary Table 3** and **Supplementary Table 4)**. We also found that out of the 3,798,220 human CCDS nullomers of length 12bp, 24 are human-specific as they were not found in any of the other 29 species queried.

Our analysis also allowed us to annotate class and order specific nullomers. For example, we found 22,738 primate-specific nullomers of 12bp length throughout the entire genome. Interestingly, this number was very close when examining 15bp primate-specific nullomers (**Figure 4c**). For nullomers of K=12bp, we observed that the majority of annotated nullomers were from non-mammalian genomes (∼96.2%), while within the mammalian nullomers, most of them were shared with primates (82%) (**Figure 4d**). We observed that humans and chimpanzees share 59% of the K=10 CCDS nullomers and 99.3% of the K=15 nullomers. In contrast, only 11% and 7% of the K=10 CCDS nullomers are shared between humans and mice or humans and yeast, respectively.

### Nullpeptide annotation

A complementary approach to identify CCDS nullomers is to identify amino acid sequences that do not appear in the human proteome, defined as nullpeptides. We scanned the UniProt human reference proteome database^10^ for nullpeptides of up to 7 aa in length. The shortest nullpeptides we found were 4 aa in length, totalling 207 across the human proteome. As expected, this number increased exponentially with length. For nullpeptides of lengths 5, 6 or 7aa, we identified 792,913, 55,524,544 and 1,269,204,068 nullpeptides, respectively.

### Nullpeptides are under negative selection

Similar to nullomers, we next examined whether nullpeptides are under negative selection. We reasoned that due to large differences in the frequency of amino acids in the proteome a more suitable metric to prioritize nullpeptides would be to sort nullpeptides by the median number of occurrences of all their possible permutations, rather than all possible single amino acid changes. We found that for a subset of nullpeptides, their permuted peptides are frequently occurring in the human proteome (**Figure 5a**), (**Supplementary Figure 2a-d**), suggesting targets for follow up experiments to understand the reason for their absence.

**Figure 5.**
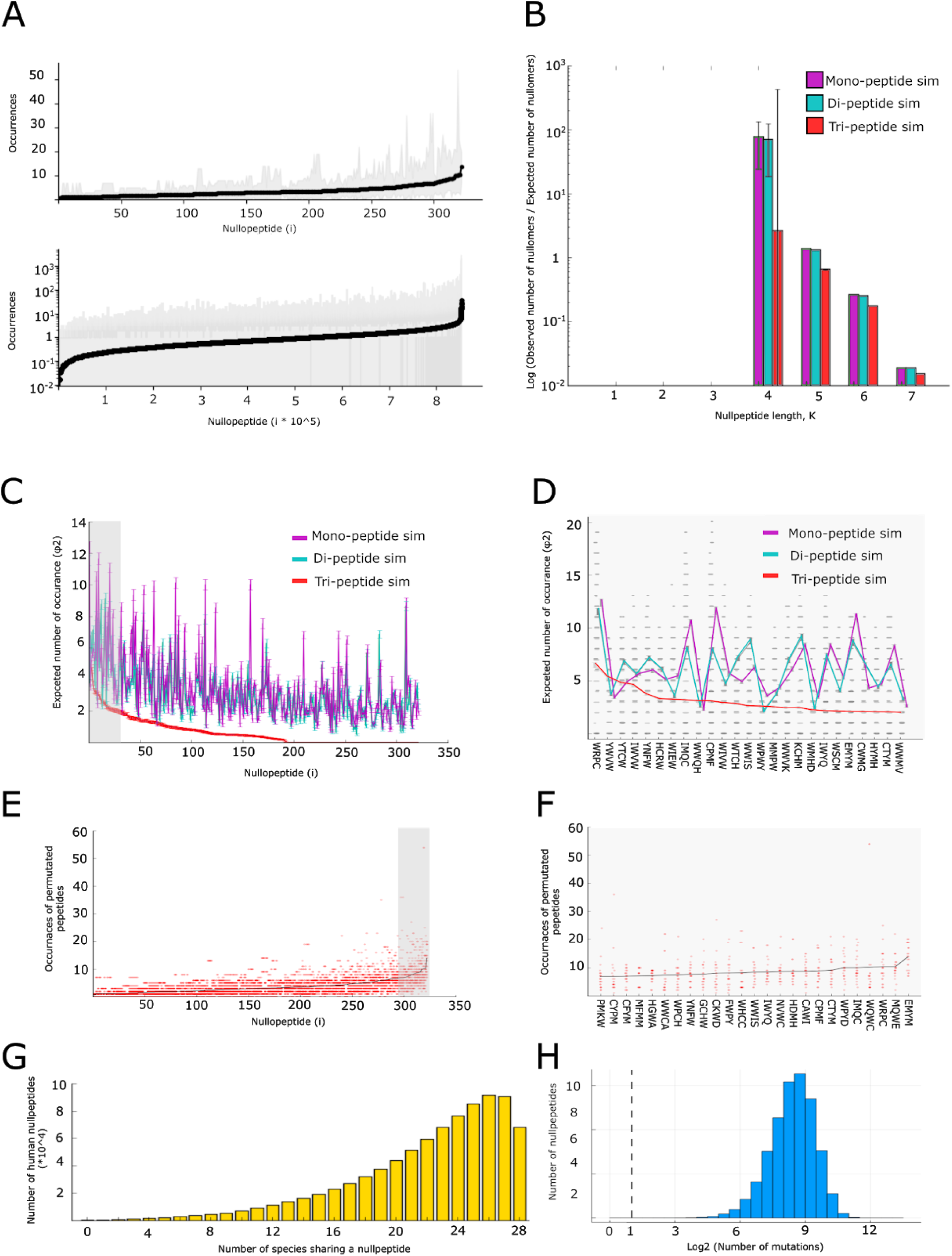
Human nullpeptide characterization. **A.** Nullpeptide prioritization based on the number of occurrences of all permuted peptides. Black line indicates the mean number of occurrences of all permuted peptides with upper and lower lines indicating minimum and maximum number of occurrences across the permuted peptides. **B**. φ2 metric score associated enrichment of nullpeptides relative to simulations controlled for the mono-, di- and tripeptide content of the proteome plotted as a function of nullomer length for K=10-15bp. **C.** Simulations showing that a large proportion of peptide nullomers (K=4) should be frequently observed in the human proteome and are likely under negative selection. Purple, turquoise and red colors represent occurrences of nullpeptides in the simulations controlling mono- di- and tri-peptide content, respectively. **D.** Depiction of the twenty five most frequent nullomers in the simulations. E. Number of occurrences of permuted peptides for each nullpeptide for K=4 aa. Black line represents the median occurrences across the permuted peptides. **F.** Twenty five top ranked nullpeptides of 4 amino acids length based on permutations. **G.** Number of species in which human nullpeptides of 4-5 amino acids length were identified. H. Distribution of nullpeptides across the number of all possible substitutions that destroy a nullpeptide.

As a second measure of selection against nullpeptides, we carried out simulations for each protein sequence of the reference proteome, controlling for monopeptide, dipeptide and tripeptide content (100-fold simulations for each). We compared the expected number of nullpeptides across the simulations and found that in all cases we observed a larger number of nullpeptides in the reference proteome than in the simulations (**Figure 5b**; **Supplementary Figure 3a-b**), suggesting negative selection against them. The expected to the observed nullpeptide frequencies showed an enrichment consistent with the results at the DNA level (**Supplementary Figure 3a-b**). For 4 aa the enrichments were 1.96-fold, 1.84-fold and 1.22-fold relative to mono- di- and tri-peptide controls and for 5 aa the enrichments were 1.19-fold, 1.14-fold and 1.08-fold relative to mono- di- and tri-peptide controls (**Supplementary Figure 3a-b**). We compared the frequency with which we observed each nullpeptide in the simulations and found that the majority of nullpeptides appeared in only a small subset of the simulations even when controlling for trinucleotide content (**Figure 5c-f; Supplementary Figure 3c-d**). Importantly, we observed consistency between our two first metrics, with nullpeptides that scored as most likely under selection in one metric also having a similar score in our second metric. As an example, among the top 25 four amino acid nullpeptides, multiple sequences were shared between the two metrics including EMYM, WRPC, IMQC, CTYM, CPMF, IWYQ, WWIS and YNFW (Wilcoxon Signed-Rank, p-value< 0.005 for top 25 nullpeptides for monopeptide, dipeptide and tripeptide comparisons) (**Figure 5d,f**). When analyzing the amino acid composition of the annotated nullpeptides, we identified the amino acids Tryptophan (W), Methionine (M) and Cysteine (C) to appear much more in nullpeptides than in non-nullomeric amino acid k-mers of the same length (**Supplementary Figure 4a**). This might be due to the importance of the M codon for translation initiation and that a single base pair change can cause W and C codons to become a TAA stop codon, the second most frequent stop codon in humans^14,15^. Similarly to nullomers, the third metric we used was the number of species in which we identified each human nullpeptide in their proteome (**Figure 5g, Supplementary Table 2**). Finally, we were also able to generate proteome-wide maps of all possible mutations that can materialize a nullpeptide and found substantial differences in the number of mutations that can generate each nullpeptide (**Figure 5h**). Combined, these results show that nullpeptides are likely under negative selection and can also be used to rank sequences for follow up functional assays.

### Characterization of nullpeptides across evolution

We next characterized nullpeptides in 28 species where we previously characterized nullomers, excluding mouse lemur for which a reference proteome was not available in UniProT (**Supplementary Tables 2-3**). We observed a negative correlation between proteome size and number of nullpeptides (**Figure 6a**) (Pearson r = −0.97). We also saw that the longer the nullpeptide, the larger the proportion was shared between species (**Figure 6b**). For 4aa, we did not find any shared nullpeptides between species. We annotated the number of nullpeptides that are only in humans but not in any other species studied, and found none for K=4aa, 135 for K=5aa and 33 for K=6aa (**Supplementary Table 5**). When expanding to primate-specific nullpeptides, i.e. missing from all primates proteomes, but could be found in at least one non-primate proteome, we found 105 4aa long nullpeptides, 23,889 for K=5aa and 7,044 for K=6aa. We believe that the drop in either human- or primate-specific nullpeptides is a result of the sharp increase in shared nullpeptides from only 5.4% of 4aa nullpeptide shared between primates and non-primates as opposed to 95.3% of the K=6aa (**Figure 6d**). We later sliced the nullpepetide lists by converging groups: humans, primates, mammals and the overall null peptide space. We observed that primate nullomers are increasing from 74.5% of mammalian nullpeptides at K=4aa, to ∼88% in K=5aa, while taking a larger portion of the total nullpeptide space at the higher K (56.4%, **Figure 6d**).

**Figure 6.**
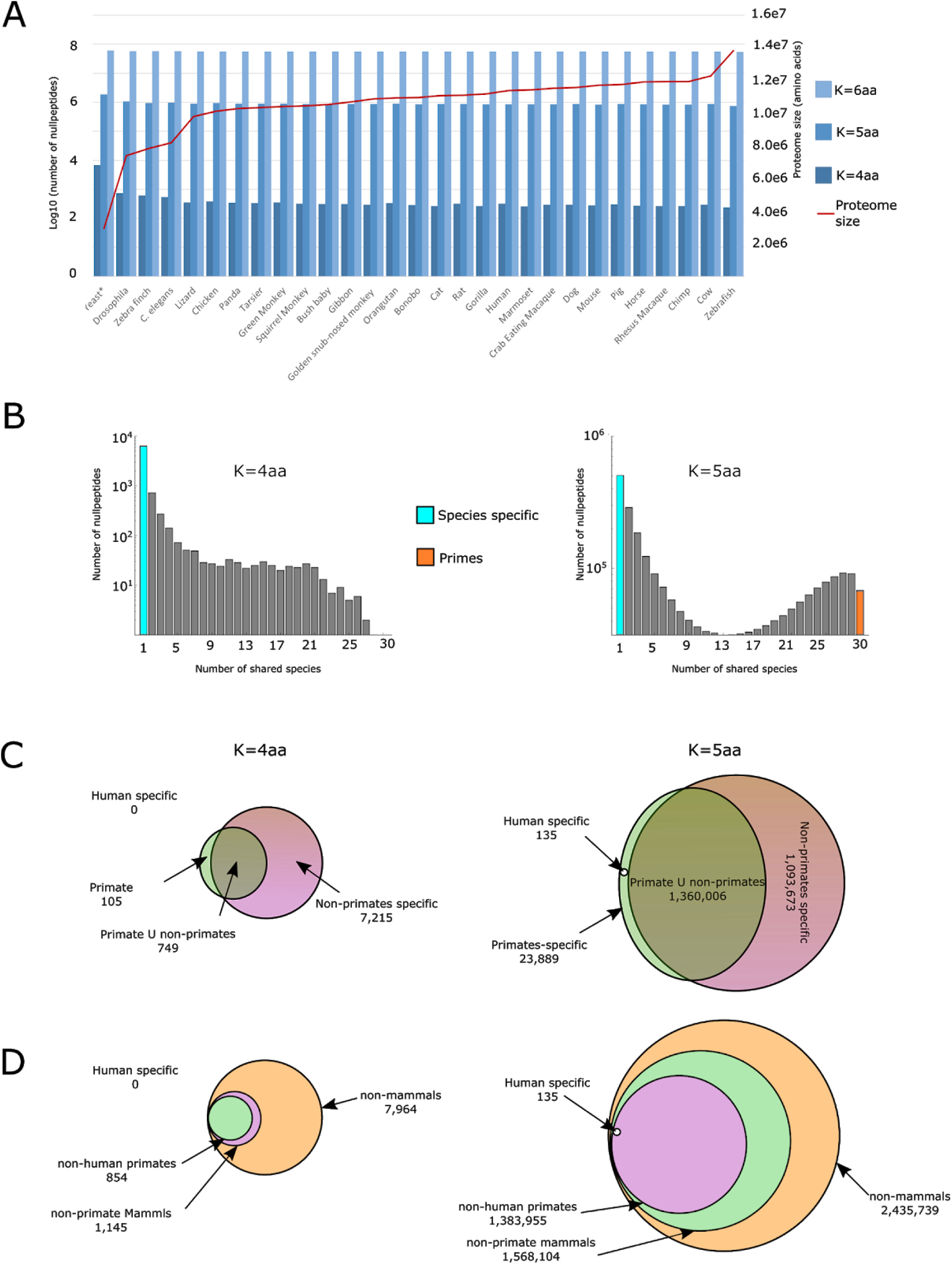
Characterization of nullpeptides across species. **A.** A bar plot displaying the number of nullpeptides found per species as a function of peptide length. Bar plot colors represent the nullpeptide lengths. The red line indicates the association between the number of nullpeptides and proteome size. **B.** Number of species sharing nullpepetides for K=4 and K=5 amino acids. Number of species-specific nullpeptides are colored in turquoise and nullpeptides shared across species are shown in orange. **C**. Venn diagrams displaying the number of nullpeptides shared between humans, other primates and non-primates species used. **D**. Venn diagrams displaying the number of nullpeptides shared between humans, all other primates aside from humans, all mammals beside primates and all species. The diameter of the circles is roughly correlated to differences in numbers.

We also annotated amino acid sequences that are absent from all known species (not just the 29 analyzed proteomes) using the UniParc database (which had 1,030,456,800 proteins), termed nullpeptide primes. We found a total of 140,308,851 nullpeptide primes, with 36,081 and 140,272,770 for six and seven amino acids in length, respectively. No nullpeptide primes were observed for K<6. To measure which amino acids are more frequent in nullpeptides, we calculated the frequency of each amino acid across prime and non-prime sequences from which we calculated the enrichment patterns. We found that the amino acids W, M, C, similar to nullpeptides, along with Tyrosine (Y) and Histidine (H) to be enriched in primes relative to non-primes, with C and W showing the strongest relative enrichment (**Supplementary Figure 4b**).

### Absent k-mers in DNA or proteome can serve as a phylogenetic classifier

Phylogenetic trees are usually built based on similarities and differences of existing sequences. Here, we wanted to test whether these trees could be built in an appropriate manner based on the absence of DNA sequences, i.e. nullomers and nullpeptides. We thus utilized our annotated nullomers and nullpeptides from the 30 species to build phylogenetic trees. We obtained an overall expected tree structure which accordingly clustered together primates, mammals and all the other organisms (**Figure 7**). The phylogenetic tree greatly improved with the length of the nullomer used. For example, we observed a significant improvement for nullomers K=14 compared to K=12 and nullpepotides K=5aa versus K=4aa, likely due to having increased numbers of sequences.

**Figure 7.**
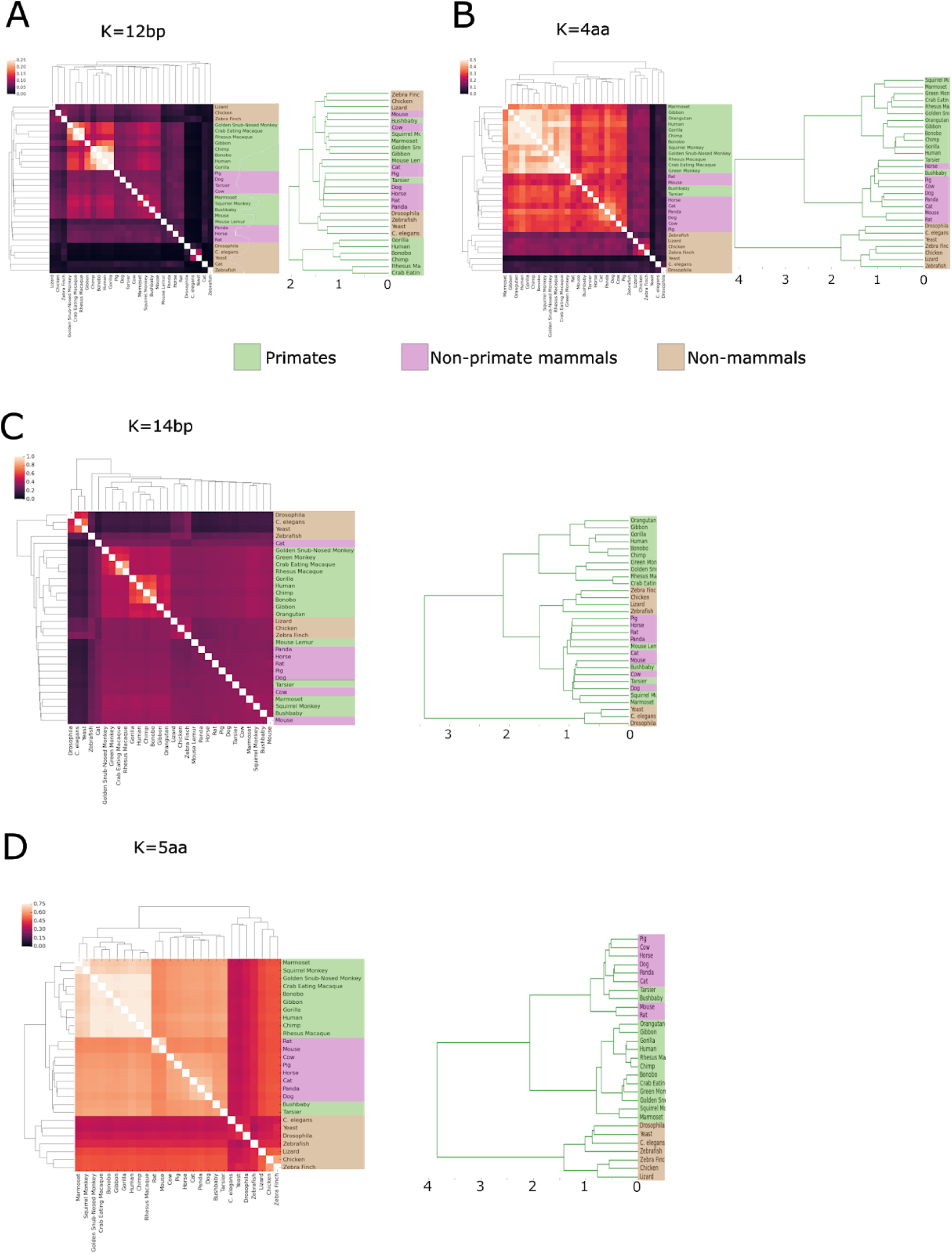
Evolutionary relationship of nullomers and nullpeptides across 30 species. **A.** Hierarchical clustering displaying the Jaccard index based on genome nullomers shared between pairs of species and the associated phylogenetic dendrogram for K=12bp displaying the evolutionary relationships between species using nullomers. **B.** Hierarchical clustering displaying the Jaccard index of peptide nullomers between species and their derived phylogenetic dendrogram for K=4aa. **C.** Hierarchical clustering displaying the Jaccard index based on genome nullomers shared between pairs of species and the associated phylogenetic dendrogram for K=14bp displaying the evolutionary relationships between species using nullomers. **D.** Hierarchical clustering displaying the Jaccard index of peptide nullomers between species and their derived phylogenetic dendrogram for K=5aa. In **A**-**D**, colors in the dendrograms represent primate (light green), non-primate mammals (light red) and non-mammals (brown).

## Discussion

Nullomers and nullpeptides are intriguing sequences whose absence in the genome or proteome could be due to their deleterious effect on the organism. Here, we characterized these sequences in the human genome/proteome and in specific functional categories. Using different approaches, we show that nullomers are under negative selection. We show that interindividual nucleotide variation could lead to the materialization of nullomers. In addition, utilizing an additional 29 genomes and proteomes from various species, we annotated both nullomers and nullpeptides that are shared among clades (primates vs. non-primates) and ones that are unique to each species (i.e. human-specific). We show that this annotation can be used to build phylogenetic trees that are similar to those using existing sequences.

To date, only three nullpeptides: WCMNW, NWMWC and WFMHW, have been tested for their effect on cell growth and apoptosis, and it was shown that the former two had an impact on cells^4,5^. In our analyses, these three sequences were all absent from the 30 species analysis, but were found 100, 93 and 241 times in the UniParc dataset from other organisms. Additionally we found that the WCMNW pentapeptide could be generated only by four substitutions across the whole proteome as opposed to an average of 400 possible mutations per nullpeptide. Utilizing our current repository and scoring criteria and expanding it in terms of nullomer length, species, genomic functional subsets etc, could assist in identifying nullomers and nullpeptides that would have a higher chance of having an effect on organismal fitness.

The use of interindividual variation to identify variant-associated nullomers could be extremely beneficial for DNA ‘fingerprinting’ purposes. These nullomers could be used to identify specific individuals or as population specific markers, as they could serve as highly sensitive probes. Probing for a specific or several specific short sequences that should be unique to an individual or population could be more straightforward and rapid both technically and computationally. A similar application could also be used to rapidly detect pathogen specific nullomers/nullpeptides. In addition, previous work has suggested that nullomers can be used to label specific DNA samples^16^. Having a list of nullomers in different functional categories and across thirty different species can greatly assist their use for sample labelling.

Phylogenetic trees are routinely constructed using existing DNA sequences, for example 16S. However, these can be complicated to construct due to horizontal/lateral gene transfer, the transfer of genetic material between unrelated organisms^17^ The use of sequences that are absent in some genomes might facilitate tree construction. While the transfer of sequences between organisms will also affect the genetic makeup of sequences that are absent, i.e. nullomers and nullpeptides, this effect might be less pronounced and as such sequence absence might pose as a useful tool to build these trees. Further analyses are needed in order to determine the advantages provided by nullomers and nullpeptides for these studies.

In summary, our work provides a list of missing DNA and amino acid sequences in over 30 genomes and in seven different functional categories. It also provides various scoring metrics to rank their potential deleteriousness effect on the organism. It will be interesting to check the functional effects of these sequences in these various categories. While coding nullomers may be more straightforward, it will be extremely intriguing to decipher the function of these sequences in the noncoding space. Past work has analyzed the occurrence of 7bp sequences in 11,257 whole human genomes to identify constrained noncoding regions and shows that they are enriched for pathogenic variation^18^. While there are no existing 7bp nullomers, it would be interesting to see whether these constrained sequences are also more enriched for nullomers and if so, how might these missing sequences affect their function.

