## Supplemental Figures 1-4 for "Absent from DNA and protein: genomic characterization of nullomers and nullpeptides across functional categories and evolution"

### Supplementary Material

**A.**

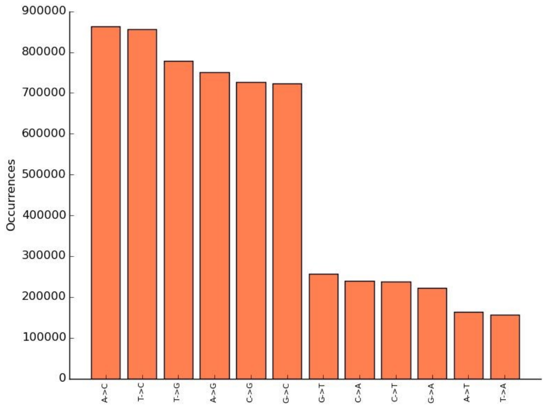


**B.**

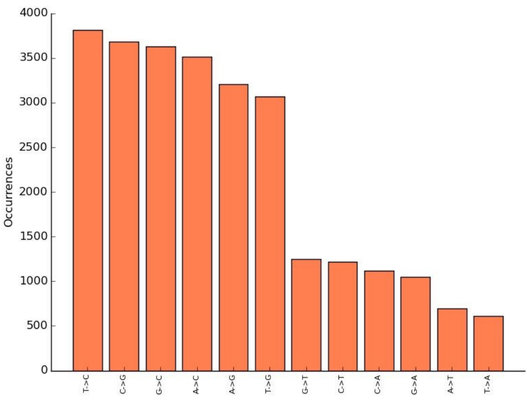


**C.**

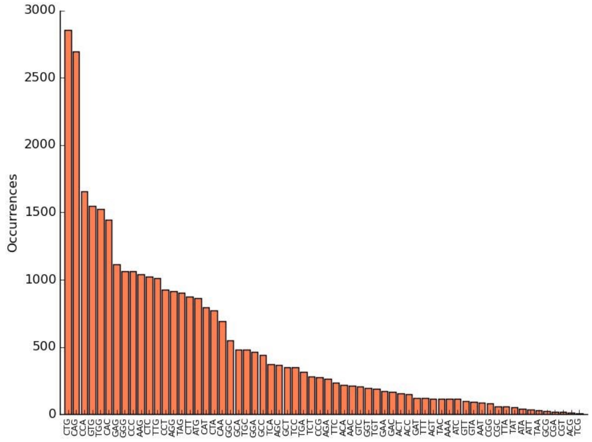


**D.**

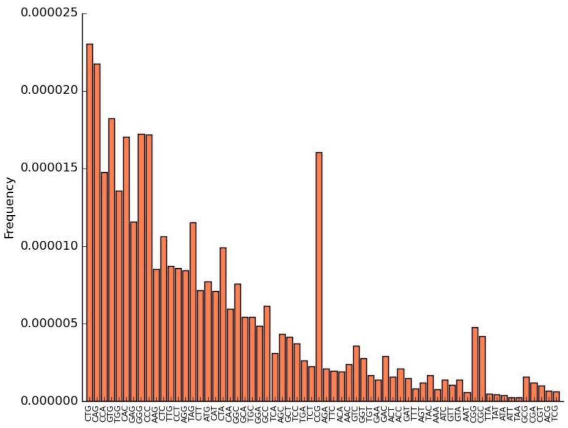


**E.**

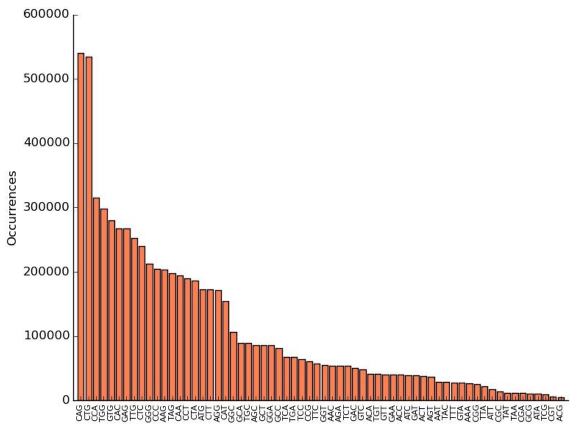


**F.**

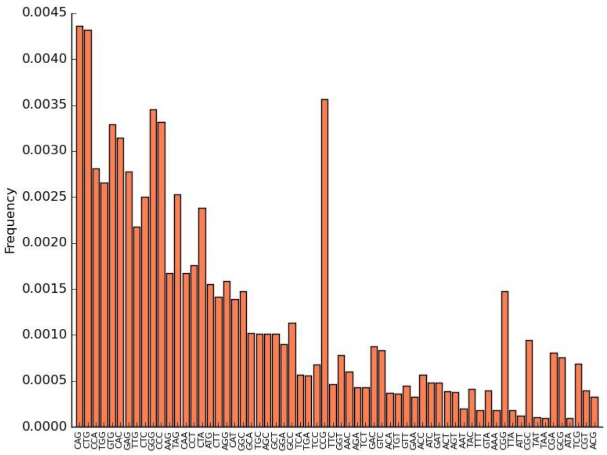

**Supplementary Figure 1. Nucleotide patterns at mutation sites that lead to the materialization of nullomers in the human genome. A.** Mutation type for substitutions for nullomer length K=11. **B.** Mutation type for substitutions for nullomer length K=12. **C.** Trinucleotide context of substitutions for nullomer length of K=11. **D.** Trinucleotide context of substitutions for nullomer length K=11, controlling for trinucleotide composition of the human genome. **E.** Trinucleotide context of substitutions for nullomer length K=11. **F.** Trinucleotide context of substitutions for nullomer length K=11 controlling for trinucleotide composition of the human genome.

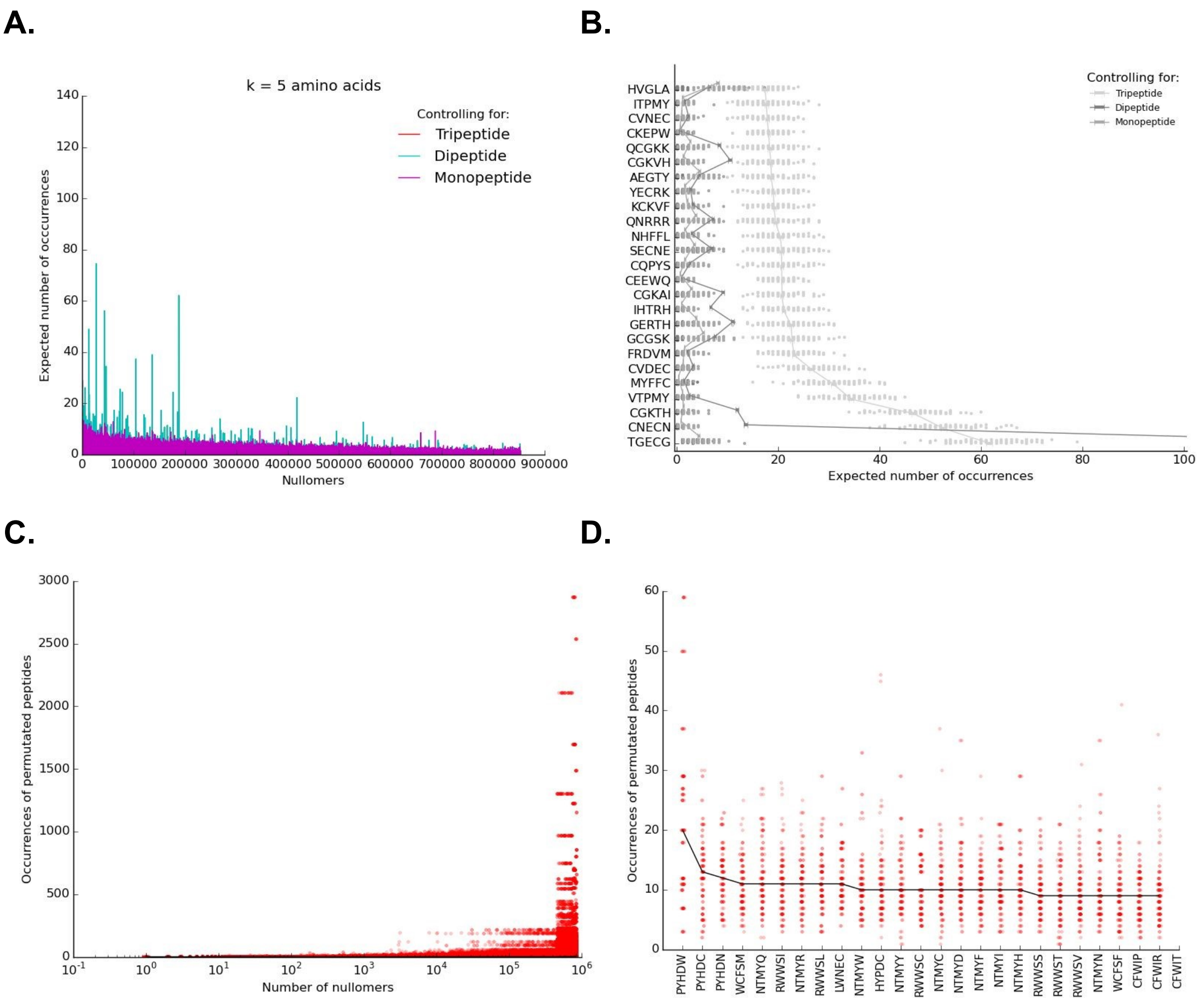


**Supplementary Figure 2. Frequency of permuted peptides for each nullpeptide in the human proteome and occurrences of nullpeptides in simulated proteomes for K=5 aa. A.** Number of occurrences of all nullpeptides in simulated proteomes controlling for protein monopeptide, dipeptide or tripeptide content for 5 aa nullpeptide length. **B.** Top 25 nullpeptides from the simulations for 5 aa length. **C.** Number of occurrences of each permuted peptide for each nullpeptide in red. The black line indicates the median occurrences for 5 aa length. **D.** The top 25 nullpeptides from the permutations for 5 aa length.

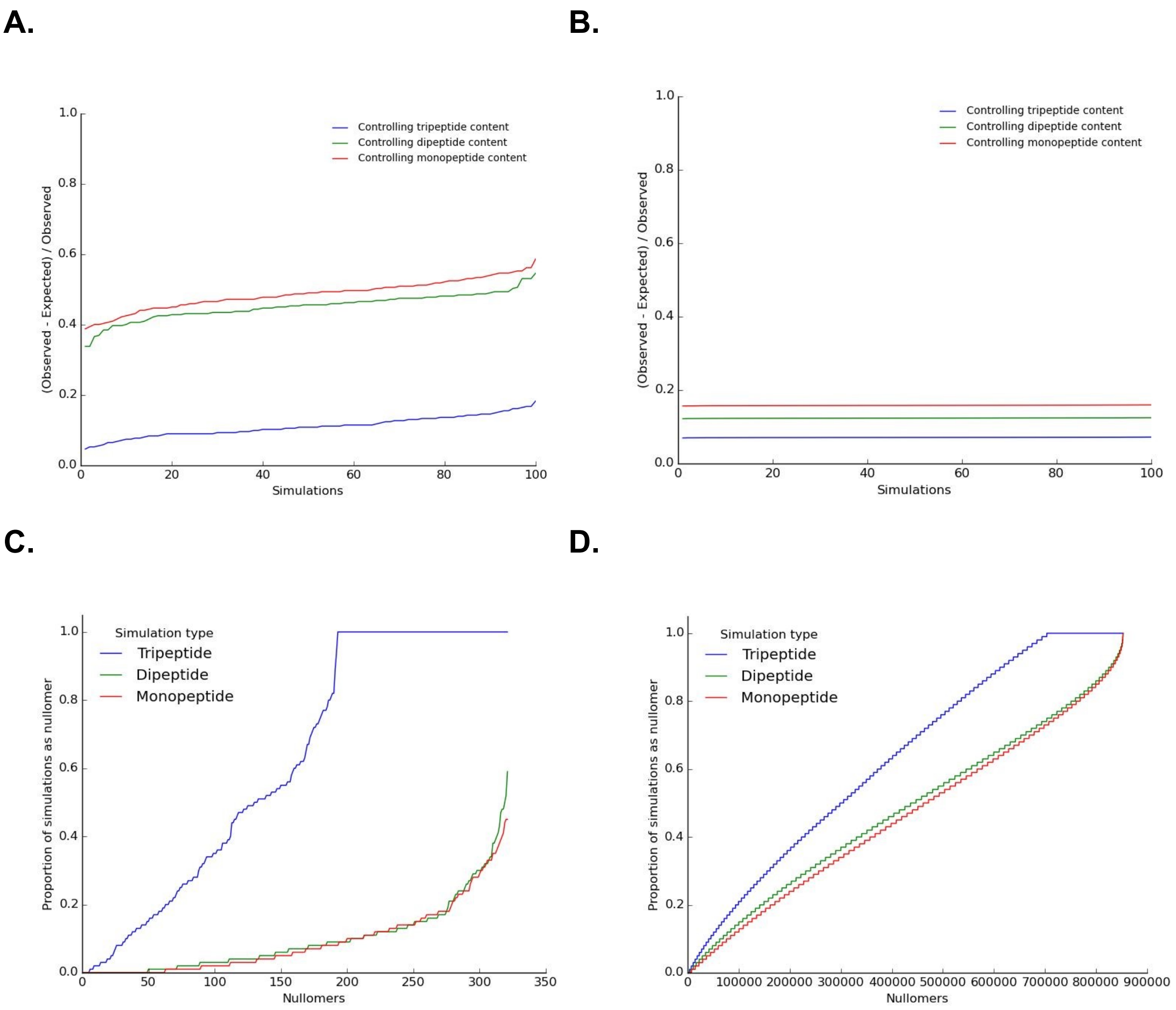


**Supplementary Figure 3. Nullpeptides are enriched in the human proteome relative to simulated proteomes.** Ratio of (Observed-Expected) / (Expected) nullpeptides across the simulations for **A.** 4 aa and **B.** 5 aa. Proportion of simulations in which each of the nullpeptides was observed for **C.** 4 aa. **D.** 5 aa.

**A.** **B.**

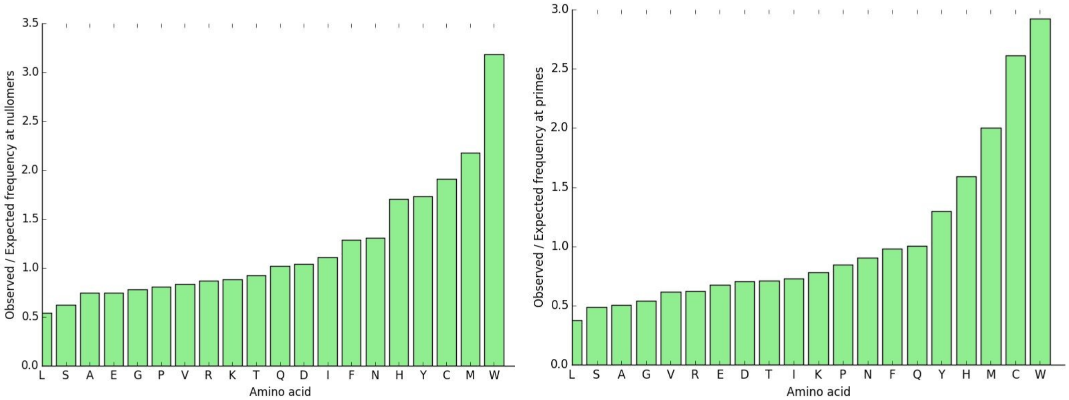


**Supplementary Figure 4. Human proteome nullpeptides and nullpeptide primes and their characteristics at the protein level. A.** Enrichment of amino acids in nullpeptide sequences compared to their frequency in non-nullpeptide sequences. **B.** Enrichment of amino acids in prime sequences and non-prime sequences from UniParc database.
