## Supplemental Tables 1-5 for "Absent from DNA and protein: genomic characterization of nullomers and nullpeptides across functional categories and evolution"

**Supplementary Table 1. Genomic nullomers per species for 30 species studied (for yeast we also found 2 nullomers at k=9).**

| Species | 10bp | 11bp | 12bp | 13bp | 14bp | 15bp |
|---|---|---|---|---|---|---|
| Zebrafish | 0 | 0 | 13,563 | 2,247,404 | 56,033,175 | 553,576,304 |
| Zebra finch | 0 | 2,330 | 278,593 | 6,920,524 | 81,100,115 | 577,652,118 |
| Yeast* | 6,455 | 463,242 | 7,229,055 | 51,021,594 | 248,244,629 | 1,051,789,596 |
| Horse | 0 | 8 | 14,351 | 1,334,788 | 31,369,815 | 363,943,354 |
| Lizard | 1 | 302 | 86,586 | 3,797,366 | 60,646,015 | 517,343,130 |
| Human | 0 | 104 | 44,287 | 2,347,664 | 40,798,250 | 405,373,474 |
| Mouse | 0 | 188 | 53,250 | 2,571,102 | 42,603,459 | 414,029,014 |
| Rat | 0 | 14 | 16,957 | 1,544,888 | 33,754,463 | 374,671,356 |
| Dog | 0 | 40 | 25,063 | 1,864,582 | 38,445,903 | 403,200,592 |
| Cat | 0 | 14 | 4,486 | 868,562 | 27,711,095 | 355,007,006 |
| Drosophila | 0 | 116 | 179,270 | 11,452,468 | 147,059,456 | 895,139,538 |
| Cow | 0 | 60 | 33,782 | 2,109,008 | 40,528,893 | 411,629,288 |
| Pig | 0 | 22 | 20,908 | 1,640,854 | 36,044,123 | 392,976,276 |
| C. elegans | 2 | 7,676 | 1,151,666 | 23,336,510 | 184,423,796 | 952,542,740 |
| Chimp | 0 | 16 | 23,740 | 1,868,770 | 37,430,940 | 392,473,026 |
| Gorilla | 0 | 142 | 45,274 | 2,388,562 | 41,156,935 | 407,034,616 |
| Bonobo | 0 | 104 | 43,560 | 2,402,008 | 41,704,635 | 410,320,644 |
| Mouse Lemur | 0 | 0 | 9,430 | 1,269,306 | 33,221,035 | 383,376,256 |
| Bush baby | 0 | 156 | 48,799 | 2,462,194 | 41,972,351 | 412,672,276 |
| Panda | 0 | 14 | 11,501 | 1,205,414 | 30,702,215 | 366,507,628 |
| Marmoset | 0 | 66 | 36,680 | 2,208,726 | 40,222,690 | 405,020,324 |
| Chicken | 0 | 250 | 88,584 | 3,981,250 | 63,134,143 | 532,125,988 |
| Gibbon | 0 | 204 | 48,375 | 2,460,574 | 41,975,895 | 411,567,194 |
| Green Monkey | 0 | 102 | 38,636 | 2,182,168 | 39,195,006 | 398,168,556 |
| Crab Eating Macaque | 0 | 110 | 39,023 | 2,193,654 | 39,327,653 | 398,895,678 |
| Orangutan | 0 | 150 | 47,633 | 2,424,730 | 41,310,233 | 406,909,060 |
| Tarsier | 0 | 0 | 24,633 | 1,771,750 | 35,420,091 | 380,837,134 |
| Golden snub-nosed monkey | 0 | 154 | 43,144 | 2,308,434 | 40,299,401 | 402,662,390 |
| Rhesus Macaque | 0 | 54 | 27,932 | 1,946,204 | 37,533,757 | 391,770,466 |
| Squirrel Monkey | 0 | 116 | 41,114 | 2,248,684 | 40,550,273 | 407,525,610 |

**Supplementary Table 2. Proteome nullpeptides per species for 30 species studied.** This list includes only the twenty standard amino-acids. No nullpeptides were observed for 3 aa in any species.

| Species | 4aa | 5aa | 6aa |
|---|---|---|---|
| Zebrafish | 237 | 751,902 | 55,170,716 |
| Zebra finch | 616 | 953,712 | 58,184,698 |
| Yeast* | 6,761 | 1,884,791 | 61,521,414 |
| Horse | 276 | 844,485 | 55,932,754 |
| Lizard | 360 | 907,843 | 56,862,023 |
| Human | 322 | 852,869 | 56,064,452 |
| Mouse | 277 | 837,543 | 55,937,143 |
| Rat | 322 | 866,503 | 56,233,869 |
| Dog | 296 | 865,856 | 56,204,050 |
| Cat | 265 | 864,460 | 56,143,883 |
| Drosophila | 737 | 1,077,673 | 58,077,699 |
| Cow | 295 | 864,760 | 56,148,113 |
| Pig | 301 | 875,880 | 56,240,045 |
| C. elegans | 543 | 979,230 | 57,533,512 |
| Chimp | 267 | 833,208 | 55,911,611 |
| Gorilla | 265 | 852,989 | 56,118,565 |
| Bonobo | 287 | 857,516 | 56,178,150 |
| Bush baby | 312 | 884,922 | 56,400,380 |
| Panda | 347 | 896,301 | 56,470,011 |
| Marmoset | 262 | 846,152 | 56,055,543 |
| Chicken | 383 | 953,712 | 57,105,793 |
| Gibbon | 314 | 872,578 | 56,331,623 |
| Green Monkey | 357 | 891,309 | 56,434,243 |
| Crab Eating Macaque | 295 | 848,487 | 56,063,665 |
| Orangutan | 337 | 901,271 | 56,564,376 |
| Tarsier | 339 | 890,858 | 56,467,613 |
| Golden snub-nosed monkey | 297 | 867,102 | 56,244,038 |
| Rhesus Macaque | 264 | 841,741 | 55,990,534 |
| Squirrel Monkey | 321 | 886,951 | 56,419,789 |

**Supplementary Table 3. CCDS nullomers per species for 30 species studied.**

| Species | 10bp | 11bp | 12bp | 13bp | 14bp | 15bp |
|---|---|---|---|---|---|---|
| Squirrel Monkey | 1,787 | 171,588 | 3,961,444 | 38,581,530 | 224,928,044 | 1,021,937,952 |
| Zebrafish | 209 | 72,814 | 2,841,598 | 34,244,162 | 216,277,759 | 1,010,397,832 |
| Zebra finch | 502,045 | 3,323,976 | 15,731,684 | 66,001,548 | 267,313,496 | 1,072,621,452 |
| Yeast | 16,828 | 680,916 | 8,538,699 | 54,316,078 | 253,176,770 | 1,057,548,108 |
| Horse | 1,655 | 171,294 | 4,075,428 | 39,388,274 | 226,889,842 | 1,024,707,006 |
| Lizard | 1,848 | 170,328 | 4,046,524 | 39,436,750 | 227,634,300 | 1,026,454,772 |
| Human | 1,553 | 159,400 | 3,798,220 | 37,728,254 | 222,865,418 | 1,018,873,404 |
| Mouse | 1,177 | 136,798 | 3,575,028 | 37,041,324 | 221,717,305 | 1,017,609,732 |
| Rat | 1,185 | 139,672 | 3,690,331 | 37,779,802 | 223,520,082 | 1,020,233,522 |
| Dog | 1,774 | 168,916 | 3,932,668 | 38,466,324 | 224,621,684 | 1,021,340,374 |
| Cat | 1,127 | 149,622 | 3,852,472 | 38,586,366 | 225,439,553 | 1,022,907,182 |
| Drosophila | 316 | 101,638 | 3,871,213 | 41,103,272 | 232,676,838 | 1,033,398,088 |
| Cow | 4,476 | 319,932 | 5,679,115 | 45,701,426 | 239,115,117 | 1,040,627,420 |
| Pig | 1,161 | 152,432 | 3,812,569 | 38,036,954 | 223,645,487 | 1,019,807,608 |
| C. elegans | 1,567 | 224,446 | 4,702,592 | 42,064,244 | 232,192,059 | 1,031,725,468 |
| Chimp | 1,398 | 152,528 | 3,707,024 | 37,296,290 | 221,914,620 | 1,017,550,102 |
| Gorilla | 1,654 | 163,094 | 3,828,991 | 37,864,388 | 223,196,164 | 1,019,391,216 |
| Bonobo | 1,636 | 164,628 | 3,857,219 | 38,010,808 | 223,559,304 | 1,019,956,808 |
| Mouse Lemur | 2,034 | 190,610 | 4,210,695 | 39,743,780 | 227,417,657 | 1,025,198,290 |
| Bush baby | 1,642 | 170,148 | 3,989,787 | 38,819,576 | 225,451,982 | 1,022,655,184 |
| Panda | 1,107 | 152,200 | 3,896,394 | 38,745,400 | 225,675,030 | 1,023,118,138 |
| Marmoset | 1,838 | 171,862 | 3,938,443 | 38,370,698 | 224,331,033 | 1,021,033,040 |
| Chicken | 13,377 | 473,650 | 5,887,884 | 43,713,660 | 233,698,888 | 1,033,256,864 |
| Gibbon | 1,840 | 171,856 | 3,945,387 | 38,444,430 | 224,561,756 | 1,021,400,348 |
| Green Monkey | 1,766 | 176,598 | 4,041,997 | 38,961,884 | 225,730,701 | 1,022,999,848 |
| Crab Eating Macaque | 1,342 | 153,716 | 3,753,448 | 37,608,610 | 222,759,700 | 1,018,874,040 |
| Orangutan | 76,771 | 1,419,604 | 11,547,865 | 60,197,684 | 260,833,866 | 1,065,940,690 |
| Tarsier | 2,671 | 212,026 | 4,417,865 | 40,597,320 | 229,217,715 | 1,027,849,846 |
| Golden snub-nosed monkey | 1,579 | 164,884 | 3,874,387 | 38,124,246 | 223,825,003 | 1,020,327,748 |
| Rhesus Macaque | 1,567 | 163,472 | 3,868,461 | 38,148,316 | 223,946,257 | 1,020,547,374 |

**Supplementary Table 4. Number of species-specific nullomers for each of the 30 species studied for K=10-15 bp.**

| Species | 10bp | 11bp | 12bp | 13bp | 14bp | 15bp |
|---|---|---|---|---|---|---|
| Squirrel Monkey | 0 | 18 | 1,480 | 2,966 | 5,194 | 1,362 |
| Zebrafish | 0 | 0 | 767 | 15,298 | 34,783 | 34,234 |
| Zebra finch | 0 | 1,270 | 39,440 | 116,668 | 105,932 | 26,488 |
| Yeast | 0 | 458,110 | 6,007,314 | 22,413,766 | 29,455,612 | 21,676,392 |
| Horse | 0 | 2 | 245 | 1,000 | 2,530 | |
| Lizard | 1 | 106 | 5,661 | 16,944 | 29,631 | 10,272 |
| Human | 0 | 24 | 994 | 1,472 | 2,470 | 454 |
| Mouse | 0 | 68 | 2,791 | 5,484 | 12,147 | |
| Rat | 0 | 4 | 380 | 1,596 | 5,000 | |
| Dog | 0 | 8 | 1,013 | 3,472 | 8,482 | 2,256 |
| Cat | 0 | 0 | 94 | 720 | 2,906 | 1,444 |
| Drosophila | 0 | 52 | 37,278 | 782,936 | 2,135,767 | 1,999,388 |
| Cow | 0 | 16 | 1,630 | 4,444 | 10,088 | 3,092 |
| Pig | 0 | 6 | 669 | 2,262 | 5,816 | 2,084 |
| C. elegans | 2 | 4,234 | 297,061 | 2,015,072 | 3,067,441 | 2,057,596 |
| Chimp | 0 | 2 | 200 | 498 | 57,176 | 174 |
| Gorilla | 0 | 20 | 1,168 | 1,738 | 2,778 | 566 |
| Bonobo | 0 | 22 | 986 | 1,352 | 1,796 | 376 |
| Mouse Lemur | 0 | 0 | 300 | 1,762 | 6,104 | 2,076 |
| Bush baby | 0 | 64 | 2,428 | 4,602 | 7,886 | 2,414 |
| Panda | 0 | 4 | 225 | 1,038 | 3,574 | 1,876 |
| Marmoset | 0 | 18 | 1,240 | 2,652 | 4,604 | |
| Chicken | 0 | 106 | 8,010 | 38,854 | 57,176 | |
| Gibbon | 0 | 74 | 1,648 | 2,712 | 4,187 | 942 |
| Green Monkey | 0 | 22 | 950 | 1,678 | 2,754 | 654 |
| Crab Eating Macaque | 0 | 2 | 627 | 878 | 1,224 | 284 |
| Orangutan | 0 | 28 | 1,549 | 2,346 | 3,704 | 766 |
| Tarsier | 0 | 12 | 753 | 1,724 | 3,662 | 898 |
| Golden snub-nosed monkey | 0 | 16 | 1,271 | 2,124 | 3,462 | 726 |
| Rhesus Macaque | 0 | 2 | 276 | 576 | 960 | 188 |

**Supplementary Table 5. Species-specific nullpeptides for each of the 29 species studied for K=4-6 aa.**

| Species | 4aa | 5aa | 6aa |
|---|---|---|---|
| Squirrel Monkey | 4 | 1,363 | 349 |
| Zebrafish | 24 | 8,377 | 4,109 |
| Zebra finch | 86 | 18,044 | 4,385 |
| Yeast | 5,559 | 342,545 | 135,006 |
| Horse | 8 | 1,440 | 390 |
| Lizard | 34 | 10,434 | 3,433 |
| Human | 0 | 135 | 33 |
| Mouse | 9 | 1,651 | 390 |
| Rat | 15 | 2,991 | 693 |
| Dog | 5 | 1,831 | 432 |
| Cat | 6 | 1,428 | 334 |
| Drosophila | 218 | 47,108 | 25,065 |
| Cow | 10 | 2,042 | 427 |
| Pig | 9 | 2,495 | 599 |
| C. elegans | 154 | 38,421 | 29,532 |
| Chimp | 0 | 178 | 53 |
| Gorilla | 2 | 675 | 257 |
| Bonobo | 2 | 527 | 174 |
| Bush baby | 19 | 2,804 | 1,006 |
| Panda | 8 | 1,925 | 488 |
| Marmoset | 5 | 1,026 | 200 |
| Chicken | 29 | 8,179 | 1,745 |
| Gibbon | 5 | 1,344 | 450 |
| Green Monkey | 10 | 1,183 | 482 |
| Crab Eating Macaque | 3 | 273 | 137 |
| Orangutan | 9 | 2,051 | 644 |
| Tarsier | 10 | 2,811 | 714 |
| Golden snub-nosed monkey | 6 | 810 | 243 |
| Rhesus Macaque | 1 | 546 | 131 |
